# Worms on the Cape: an integrative survey of polydorid infestation in wild and cultivated oysters (*Crassostrea virginica*) from Massachusetts, USA

**DOI:** 10.1101/2023.08.12.553105

**Authors:** Andrew A. Davinack, Margaret Strong, Barbara Brennessel

**Affiliations:** Biology Department, Wheaton College. 26 East Main Street, Norton, MA 02766, USA

**Keywords:** polydora, mollusca, polychaete, aquaculture, websteri

## Abstract

Polydorid infestations pose a significant challenge to shellfish aquaculture by impacting marketability and profitability of farms. In this study, we investigated the prevalence, intensity, identity and biogeography of shell-boring worms infecting both farmed and wild oysters (*Crassostrea virginica*) from three sites in Wellfleet Harbor, Massachusetts – an economically important shellfishing region in the northeastern United States. DNA barcoding revealed that *Polydora websteri* was the sole culprit responsible for infecting oysters from all three sites, reaching maximum prevalence (100% infection) and intensity (mean intensity: 38.63) in the Herring River. The oysters in the Herring River are subjected to restricted tidal flow due to the presence of a physical barrier (dike), and this could be responsible for the high infestation levels of *P. websteri* observed in this population. In addition, a population genetic analysis incorporating COI sequence data from Wellfleet *P. websteri* in addition to newly published sequences from the Black Sea and the Sea of Azov found very low levels of genetic differentiation across several intercontinental populations (0.000 – 0.399), which is likely being driven by multiple introductory events such as oyster importations. These findings are discussed in relation to the future of shellfish aquaculture in the United States.

## 1. Introduction

The shellfish aquaculture trade is a $100B industry and plays a vital role in meeting the global demand for seafood, providing a sustainable source of protein and contributing to economic growth in coastal regions (Azra et al., 2021). With the decline of wild fisheries and increasing concerns about overfishing, the cultivation of shellfish has emerged as a promising solution to meet the growing demand for marine resources (Dalton et al., 2017; Azra et al., 2021). Among the diverse range of commercially cultivated shellfish species, the eastern oyster (*Crassostrea virginica*) holds special significance due to its ecological, economic and cultural importance in coastal ecosystems and communities along the east coast of the United States (Botta et al., 2020; Beckensteiner et al., 2020). Oyster aquaculture has experienced remarkable expansion worldwide, driven by advances in hatchery techniques, improvements in husbandry practices, and increasing consumer demand for high-quality oysters. In the United States and New England in particular, some form or another of oyster aquaculture has occurred since pre-colonial times with intensive systematic husbandry and cultivation occurring shortly after the first European colonists arrived (Brennessel, 2008; Botta et al., 2020). In the small region of Cape Cod alone, there are more than 250 oyster growers spread out across 700 acres and they range from large scale operations to single person sites (Brennessel, 2008).

The success of oyster aquaculture is not without its challenges. Oyster populations, both wild and cultivated, are vulnerable to a range of biotic and abiotic stressors (Brennessel, 2008; Botta et al., 2020). Biotic factors include predation (which includes overharvesting by humans), competition for resources, and diseases. In fact, a combination of overharvesting and diseases have decimated many natural shellfish beds across Cape Cod (Brennessel, 2008). As a consequence, oyster farmers here can no longer depend on natural replenishment of shellfish stocks and are now dependent on a reliable seed supply from both local and regional shellfish hatcheries (Brennessel, 2008). One biotic stressor that has garnered increasing attention due to its negative impact on oyster growth, survival and marketability is the so-called ‘mud-blister’ worms. These are tubicolous polychaete worms of the Spionidae family (sometimes referred to as ‘polydorids’ or ‘polydorins’) which are capable of burrowing into calcareous substrates (Silverbrand et al., 2020; Davinack and Hill, 2022; Tan et al., 2023). Shell-boring polydorids are largely sessile and are dependent on their larvae for dispersal. Once the larvae are liberated from the female’s egg string, they then settle on a calcareous substrate (e.g., an oyster), undergo metamorphosis and begin burrowing into the shell where they henceforth assume a sessile existence carrying out all of their essential biological activities, including reproduction and feeding (David, 2021). The burrowing activity of the worms weakens the oyster shell and forces the oyster to divert energy from growth to shell-repair resulting in smaller sizes at maturity and the emergence of unsightly blisters on the innermost layer of the shell, both of which directly impacts marketability of the crop and therefore profitability of the farm (Williams et al., 2016). In fact, heavy infestation of polydorid worms have led to the closure of several shellfish farms around the world (Tan et al., 2023). In the United States, *Polydora websteri* has historically been recorded as the main culprit for shell infestations in commercial bivalves (Martinelli et al., 2020; David, 2021). However, in the last few years, two additional species – *Polydora onagawensis* and *Polydora neocaeca* have emerged as pathogens of eastern oysters and bay scallops, respectively, in the northeastern United States (Silverbrand et al., 2021; Davinack and Hill, 2022). Coupled with climate change, these polychaete pests pose a serious threat to to shellfish stocks throughout the region (David, 2021).

To assess the impacts of polydorid worms on their shellfish hosts, researchers have focused on studying their prevalence, distribution and identities, the latter of which has incorporated molecular approaches such as DNA barcoding (Simon and Booth, 2007; Simon et al., 2009; Davinack and Hill, 2022). In this study we used an integrative approach to assess polydorid infestation in both wild and cultivated stocks of *C. virginica* from Wellfleet Harbor – a major oyster farming region in Cape Cod, Massachusetts. This survey is particularly interesting because the wild oysters were sampled from a contaminated estuary (the Herring River system) where oyster harvesting is prohibited by governmental authorities due to high fecal coliform bacteria levels (Portnoy and Allen, 2006). The farmed oysters are located at sites which have been approved by environmental authorities for harvesting and planting of seed – thereby acting as ‘control’ sites for this study. Interestingly, all three sites are located no more than 10 km from each other.

The objectives of this study were two-fold: (1) to determine and compare the levels of polydorid infestation in ‘wild’ shellfish at the contaminated site and at the two sites where oysters are cultivated, harvested and commercially sold, and (2) to determine the identity and genetic interrelationships of these shell-boring polydorids from the three sites using DNA barcoding and population genetic analyses.

## 2. Materials and Methods

### 2.1. Brief Description of Study Sites and Farming Operation

Three localities in Wellfleet Harbor were selected for sampling: Herring River (‘wild’ stock), Mayo Beach (cultivated stock) and Black Fish Creek (cultivated stock) (Fig. 1). All three sites are subjected to high tidal fluctuations with averages of 2 – 3.5 m above mean low water at high tide. Mayo Beach (MB) is located near the town marina and has several licensed shellfish growing areas that have been cultivated since the 1980s. Black Fish Creek (BFC) is located south of the main section of Wellfleet Harbor and has relatively more connectivity to Cape Cod Bay. The Herring River system (HR) is a 1,100-acre estuary that has been tidally restricted for decades due to the construction of a dike at the mouth of the river (Portnoy and Allen, 2006). The ‘wild’ oysters located at this site are underwater during all tidal regimes and harvesting is prohibited due to high coliform bacteria levels. In Wellfleet, oysters are usually cultivated via a ‘bags on rack’ system such that oysters are grown approximately 60 cm higher than the sediment and usually exposed to air at low tide. The farmed oysters (MB and BFC populations) are usually ready for harvesting and trade after 18-24 months and are thus removed after this time, while the ‘wild’ oysters from the Herring River remain *in situ* due to harvest restrictions.

**Fig. 1.**
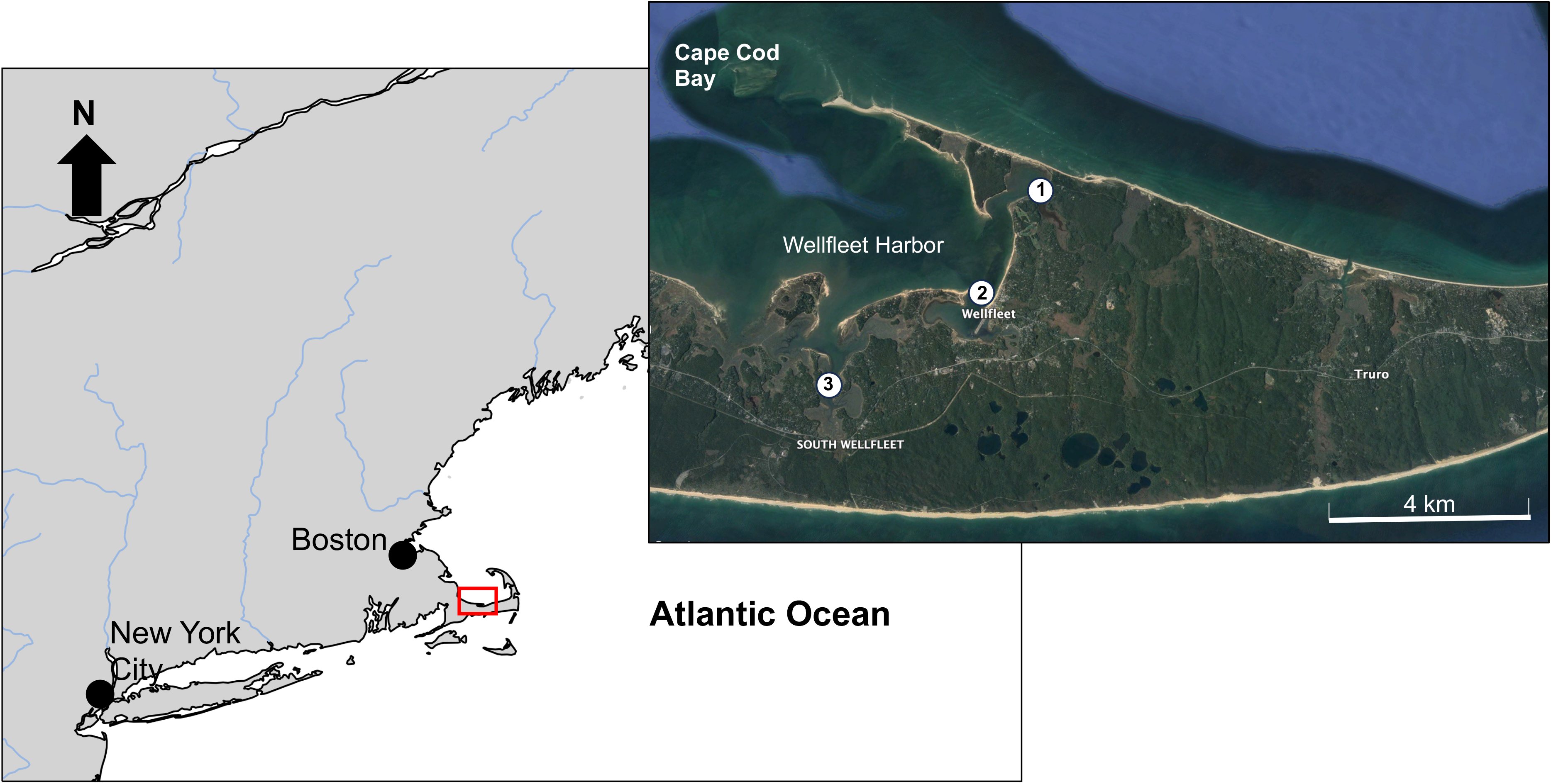
Map showing sampling localities of the eastern oyster *Crassostrea virginica* from the Cape Cod region of Massachusetts, USA. 1 = Herring River, 2 = Mayo Beach, 3 = Black Fish Creek.

### 2.2. Oyster Sampling and Worm Extraction

Thirty oysters were sampled from each of the three sites on May 30, 2023 during low tide based on the quota restrictions of our collections permit (Massachusetts DMF Special Collections Permit #185362). For the HR site, oysters were collected at random by hand with the aid of clam rakes. For the MB site oysters were supplied by farm staff during harvesting and for the BFC site, oysters were taken from different oyster racks which were stacked on top of each other. Temperature, salinity and pH measurements were recorded for each site (Table 1).

**Table 1.** Summary data of *Crassostrea virginica* sampling sites in Wellfleet, Massachusetts.

| Site | GPS<br>coordinates | Temperature<br>(°C) | Salinity<br>(psu) | pH | Cultivation<br>status | Number of<br>oysters<br>sampled (N) | Mean<br>oyster<br>size<br>± SD |
| --- | --- | --- | --- | --- | --- | --- | --- |
| Herring<br>River | 41°55'54' N<br>70°03'56' W | 21.4°C | 24.6 | 8.3 | Wild | 30 | 7.12 ±<br>1.43 |
| Mayo<br>Beach | 41°55'44' N<br>70°01'59' W | 19.9°C | 29.3 | 8.3 | Farmed | 30 | 7.61 ±<br>0.74 |
| Black<br>Fish<br>Creek | 41°54'33' N<br>69°59'51' W | 22.0°C | 30.1 | 8.2 | Farmed | 30 | 7.53 ±<br>1.73 |

Oysters were transported to a mobile lab and each individual was tagged and size measurements were taken using a vernier caliper. Each oyster was then immersed in a 500 ml beaker filled with a 0.05% phenol solution for 45 mins to flush out any worms that might be occupying the outer prismatic layer of the shell. After 45 mins, oysters were checked to determine whether worms had evacuated their burrows and any worms present were removed and placed in a glass petri dish labelled with the corresponding oyster tag. Oysters were then shucked, the meat removed and photos were taken of both shell valves. Shells were then carefully broken with pliers to minimize damage to worms. Each shell was thoroughly checked and rechecked for worm burrows and all worms were removed via forceps or pipettes and placed into their appropriate petri dishes. Live worms were then identified morphologically using the most recent regional descriptions of *Polydora* borers (Silverbrand et al., 2021; Davinack and Hill, 2022) and then placed in 99% molecular grade ethanol for genetic analysis.

### 2.3. DNA Extraction, PCR and Sequencing

Genomic DNA was extracted from six worms per site (18 specimens in total) using the D’Neasy Blood and Tissue Kit (Qiagen, Hilden, Germany) following the manufacturer’s instructions. DNA quantity ranged from 10.2 to 56.5 ng/ul. Polymerase Chain Reaction of the cytochrome c oxidase I (COI) gene was carried out using the Dorid COI.3F and Dorid COI.1R polydorid-specific primer pairs from Williams et al. (2017) along with its corresponding cycling conditions. PCR products were amplified on a 2% agarose gel and amplicons were excised and cleaned using a gel purification kit (Qiagen, Hilden, Germany). Purified DNA was sequenced using both forward and reverse primers and Big Dye Terminator Cycling Sequencing at Azenta LLC (Plainfield, NJ).

### 2.4. Data Analysis

#### 2.4.1. Infection Data

To determine parasite load, prevalence was first calculated as a percentage by dividing the number of infected hosts by the total number of hosts. Intensity was determined as the total number of polydorid worms per oyster and mean intensity was calculated for each sampling site. To determine whether there was a significant difference in mean intensity across sampling sites, a Kruskal-Wallis H test was carried out followed by a Dunn’s post-hoc test to determine where significance (if any) lay. To determine the magnitude of intensity among sites, Cohen’s *d* was also calculated in a pairwise manner. To determine whether there was a relationship between mud-blister coverage and intensity, the surface area of each shell valve per oyster along with the total surface area of the mud-blister for each valve were measured using the freeform tool in Image J (Schneider et al., 2012). The percentage of the shell with mud-blisters was then calculated for each oyster and a regression analysis was carried out between polydorid intensity and percentage shell surface with mud blisters to determine if the latter can be a potential predictor for parasite load. All statistical analyses were performed in Python 3 (Van Rossum and Drake, 2009).

#### 2.4.2. Genetic Data

Sequences were first compared against the GenBank database using the BLASTn algorithm and then all sequences were translated into amino acids using the ExPASy translation tool to check for stop codons. Preliminary BLASTn comparisons found that all 18 individuals sequenced were 99.36% to 99.84% similar to *Polydora websteri*, which corroborated initial morphological assessments in the field. All sequences were deposited into the GenBank database (accession numbers: OR430058 – OR430075). A representative dataset consisting of common polydorids associated with molluscs in the eastern United States was then assembled which included the *P. websteri* sequences from this study (Supplementary Table 1). Sequences were aligned and edited using the Clustal W alignment algorithm implemented in Bioedit ver 5.0.6 (Hall, 1999). The phylogenetic position of *P. websteri* was assessed by constructing a maximum-likelihood (ML) tree in MEGAX (Kumar et al., 2018). The GTR+G nucleotide substitution model was selected as the best fitting model, determined using the corrected Akaike Information Criterion best fit model test (AICc) in jModelTest ver. 2.0 (Darriba et al., 2012). Pairwise kimura-2-parameter (K2P) genetic distances were then calculated in MEGAX to determine intra and inter-specific genetic differences.

To determine both regional and global genetic connectivity patterns between *P. websteri* from the three Wellfleet sites and those from other geographic localities, a larger dataset was assembled consisting of all verified sequences of *P. websteri* on the GenBank database (Table 2). A TCS haplotype network was then constructed in PopART ver. 1.7 (Leigh and Bryant, 2015) to determine the evolutionary relationships among haplotypes. Finally, to estimate levels of genetic differentiation, populations were clustered based on geographic locality with specimens from the Herring River, Black Fish Creek and Mayo Beach all aggregated into a single cluster (Wellfleet) and both a hierarchal Analysis of Molecular Variance (AMOVA) and pairwise AMOVA calculations (Φ_ST_) were carried out in Arlequin ver. 3.5 (Excoffier and Lischer, 2010). Pairwise genetic differentiation estimates were only calculated for clusters which had more than five individuals per population (n > 5).

**Table 2.**
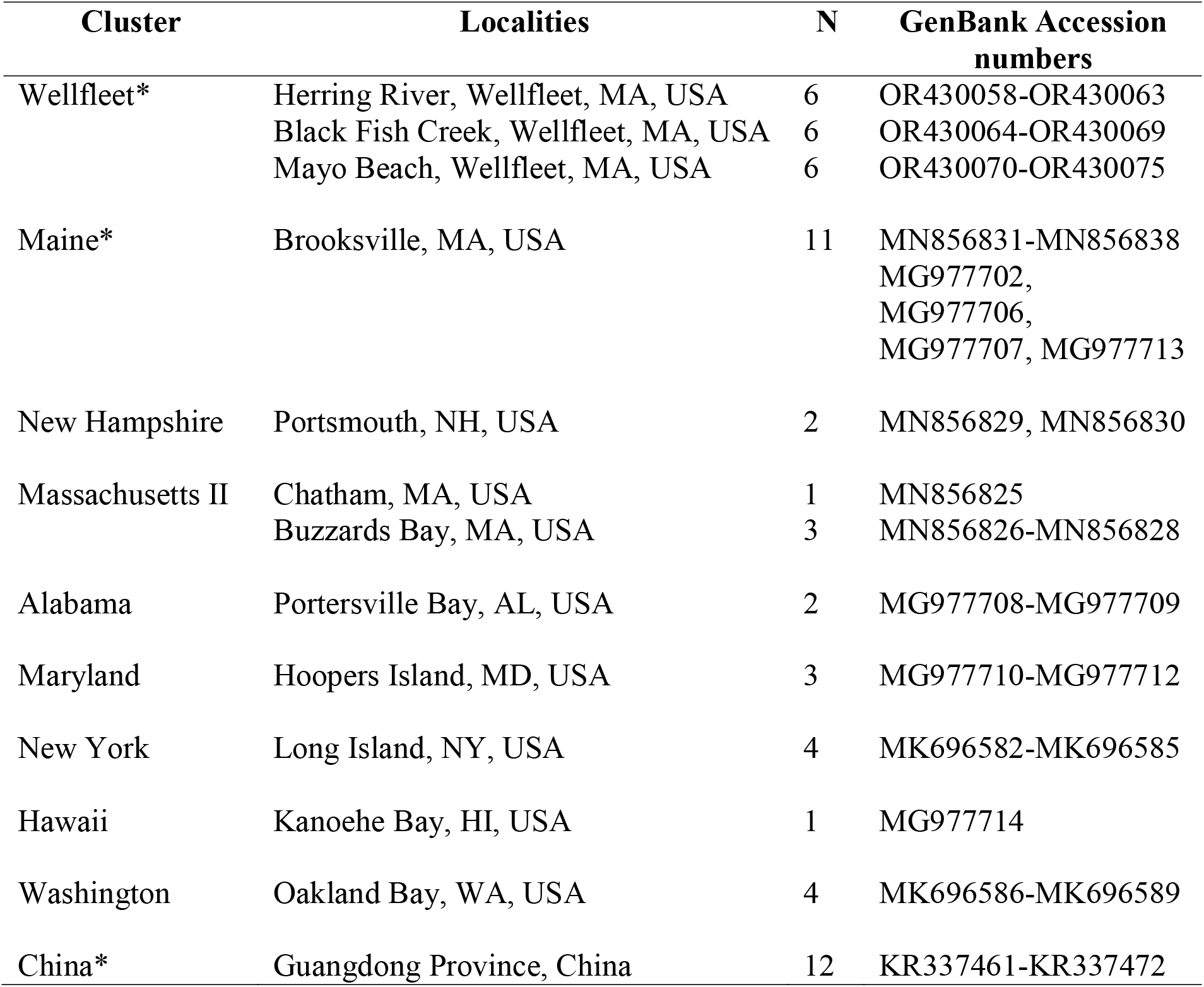
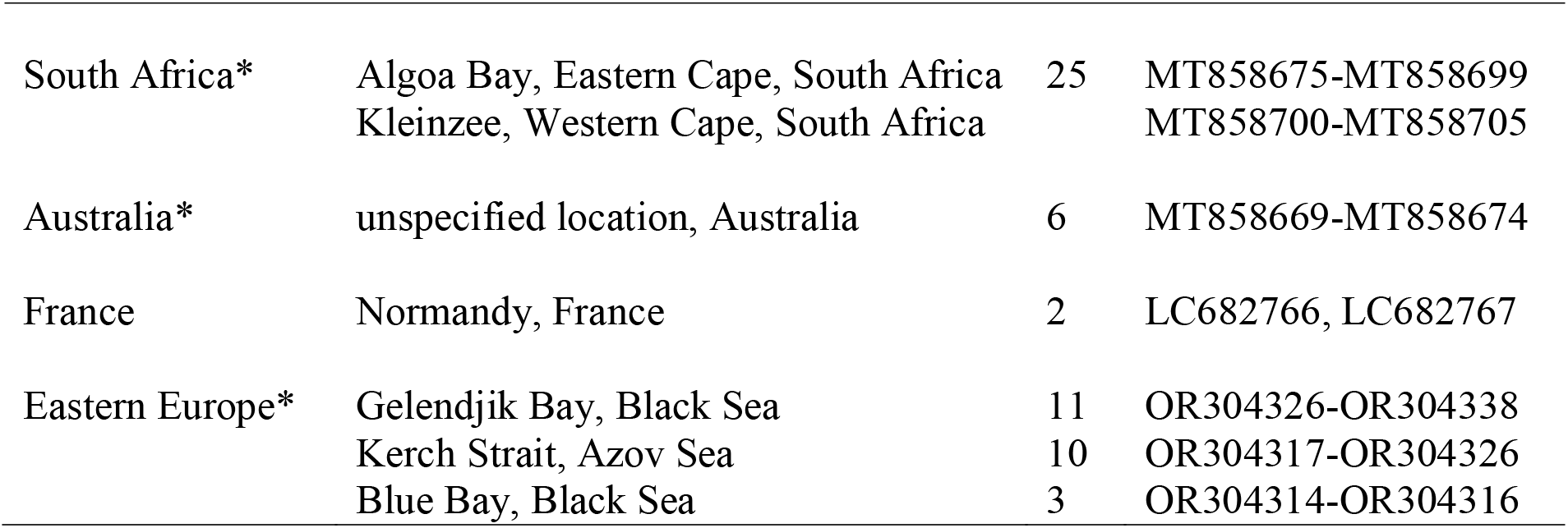
Mitochondrial COI sequence data used to assess connectivity and genetic interrelationships in *Polydora websteri*. *used in Φ_ST_ calculations.

## 3. Results

### 3.1. Genetic analyses

A maximum-likelihood phylogenetic tree nested all 18 sequenced specimens into a highly supported *P. websteri* clade with intraspecific genetic distances ranging from 0.000 to 0.011 (Fig. 2). Interspecific genetic distances with other common east coast polydorids such as *P. neocaeca*, *P. onagaewensis* and *P. cornuta* ranged from 0.164 – 0.219. A haplotype analysis recovered 22 haplotypes of which nine were shared and 13 were unique with the resulting haplotype network showing no geographic patterning of haplotypes. In fact, several *P. websteri* individuals from the Wellfleet region shared a single common haplotype with 13 other populations, some of which span multiple intercontinental ocean basins (Fig. 3). A hierarchical AMOVA corroborated this by showing that the majority of the genetic variation was found within populations (89.5%) rather than among them (10.5%). Pairwise genetic differentiation estimates were relatively low and ranged from 0.000 – 0.399. A pairwise AMOVA found that *P. websteri* from Wellfleet showed the highest levels of connectivity with worms from Maine (Φ_ST_ = 0.013) and exhibited the lowest levels of connectivity (more structuring) with worms from China (Φ_ST_ = 0.138) and South Africa (Φ_ST_ = 0.137) (Table 3).

**Fig. 2.**
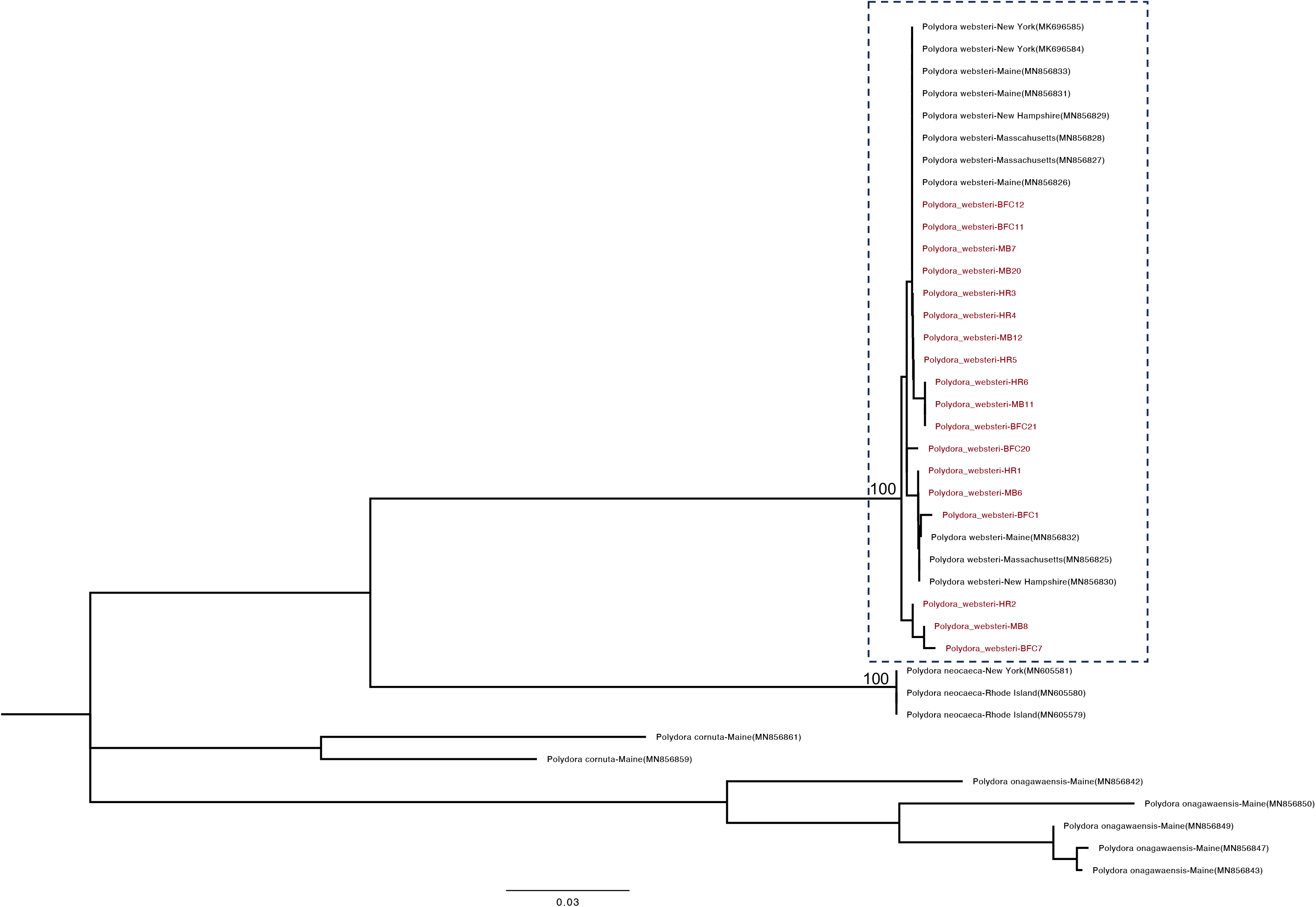
Maximum-Likelihood (ML) tree of COI barcodes showing reciprocal monophyly of *Polydora websteri* worms (in red) extracted from *Crassostrea virginica* from Wellfleet Harbor in Cape Cod. In-group taxa includes published COI barcodes of known shell-boring polydorids from the eastern United States and polydorids often associated with commercial molluscs. Values above nodes represent bootstrap support based on 1000 replications.

**Fig. 3.**
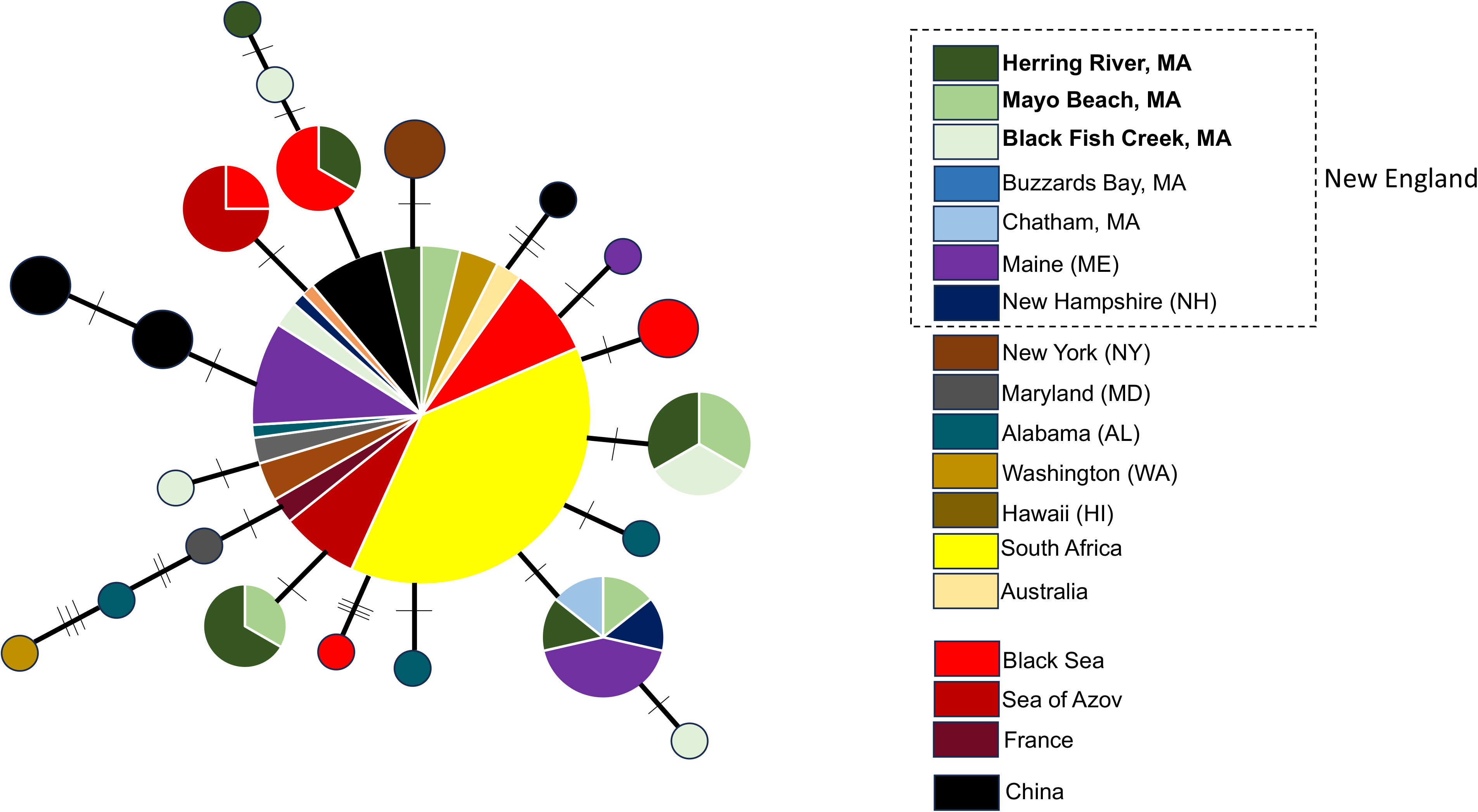
Haplotype network for *Polydora websteri* based on mtDNA – COI sequence data. Size of circles is representative of individuals with that haplotype. Smallest circles represent a haplotype frequency of one. Each connecting line between haplotypes represents one mutational step and notches on connecting lines represent additional mutational changes.

**Table 3.**
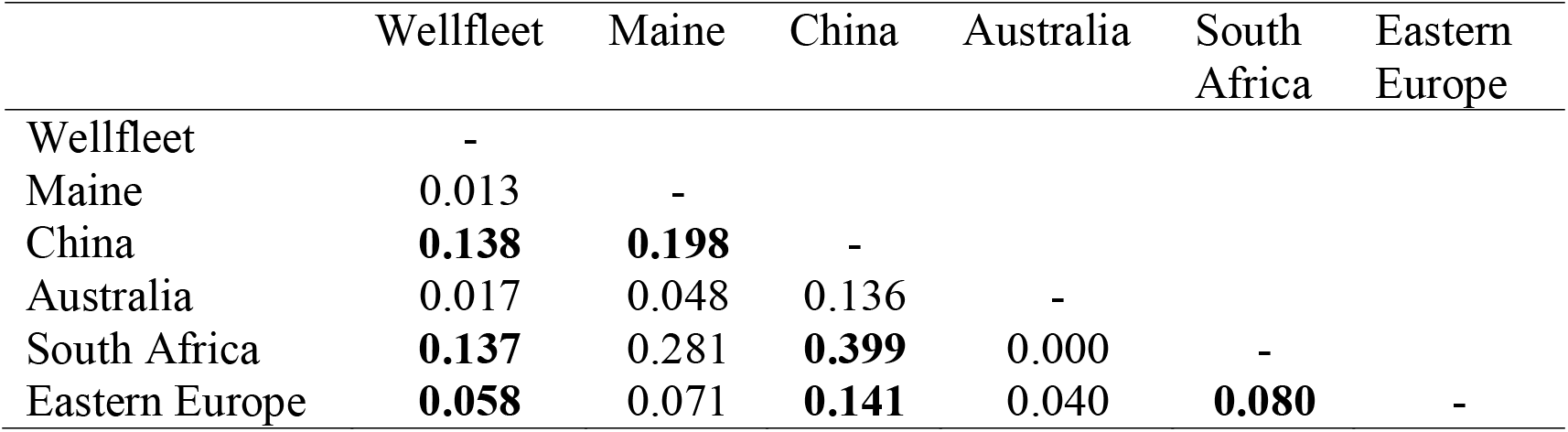
Pairwise genetic differentiation (Φ_ST_) estimates for selected *Polydora websteri* geographic clusters based on the mtDNA COI marker. Bolded values indicate significance (p < 0.05).

|  | Wellfleet | Maine | China | Australia | South Africa | Eastern Europe |
| --- | --- | --- | --- | --- | --- | --- |
| Wellfleet | - |  |  |  |  |  |
| Maine | 0.013 | - |  |  |  |  |
| China | **0.138** | **0.198** | - |  |  |  |
| Australia | 0.017 | 0.048 | 0.136 | - |  |  |
| South Africa | **0.137** | 0.281 | **0.399** | 0.000 | - |  |
| Eastern Europe | **0.058** | 0.071 | **0.141** | 0.040 | **0.080** | - |

### 3.1. Infestation data

Polydorid prevalence was highest at the Herring River (HR) site where 100% of oysters sampled were infected, while Black Fish Creek (BFC) and Mayo Beach (MB) had an overall prevalence of 83% and 73% respectively (Fig. 4). Brooding females of *P. websteri*, in addition to their characteristic three-chaetiger larvae were observed in oyster samples from all three sites (Fig. 5). Mean polydorid intensity for the HR, BFC and MB sites were 38.63 ± 17.58, 8.72 ± 4.96 and 16.84 ± 9.41 respectively (Fig. 6). There was a significant difference in polydorid intensity among the three sites (H = 48.14, p < 0.05) and a Dunn’s post-hoc test showed that polydorid intensity of oysters from HR was significantly higher than those reported from MB and BFC (p < 0.05) while polydorid intensity between MB and BFC was not significant (p = 0.06). A pairwise comparison of the magnitude of intensity differences was lower for MB and BFC (Cohen’s *d*: 0.10) than for HR and the other two sites (Cohen’s *d*: 0.19 and 0.17 for MB and BFC respectively). Finally, there was a strong positive and significant relationship between mud blister surface area and polydorid intensity (R^2^ = 0.79, F = 323.7, p < 0.05, N = 90).

**Fig. 4.**
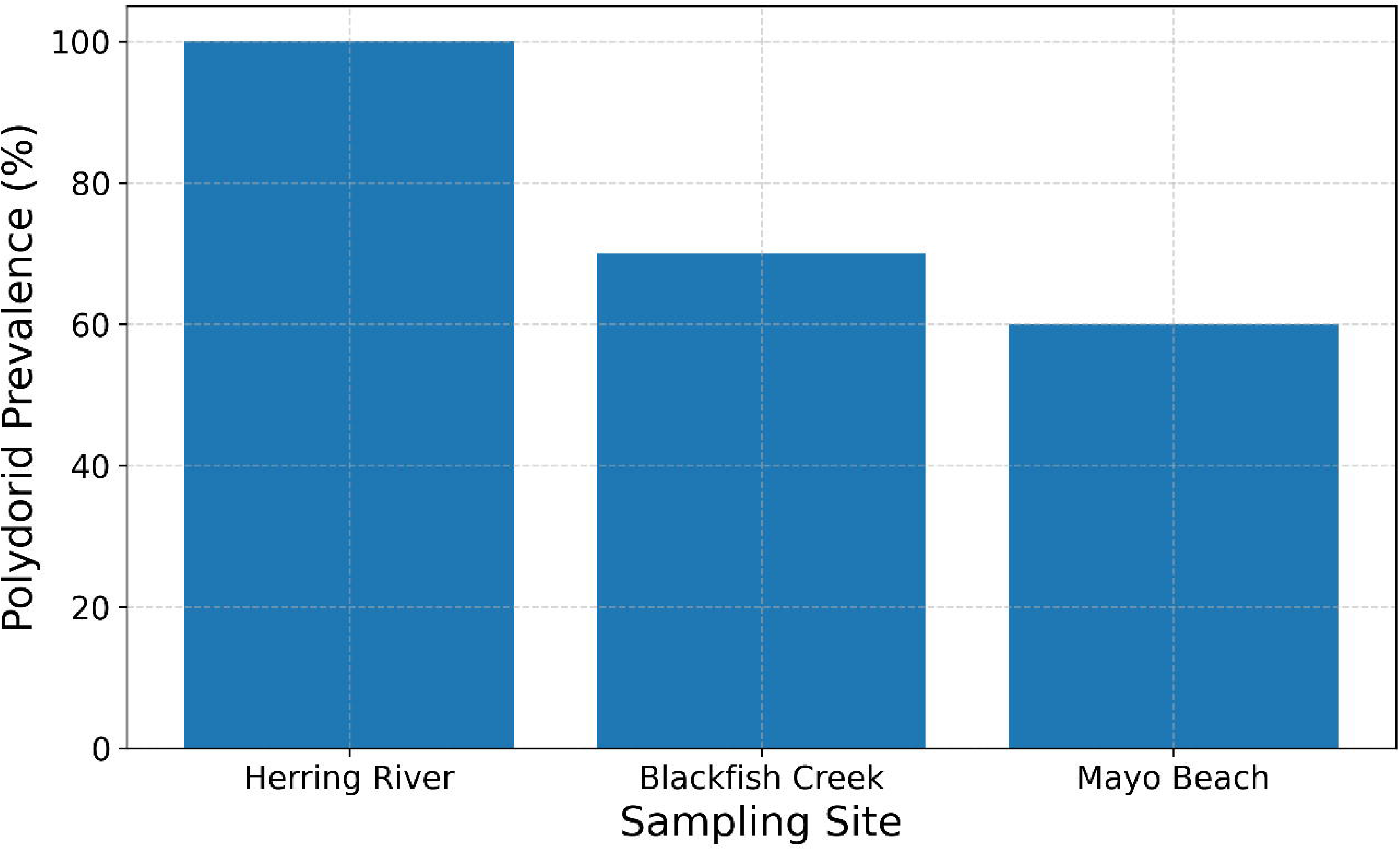
Bar graph showing prevalence of *Polydora websteri* in oysters (*Crassostrea virginica*) sampled from three sites in Wellfleet Harbor, Massachusetts.

**Fig. 5.**
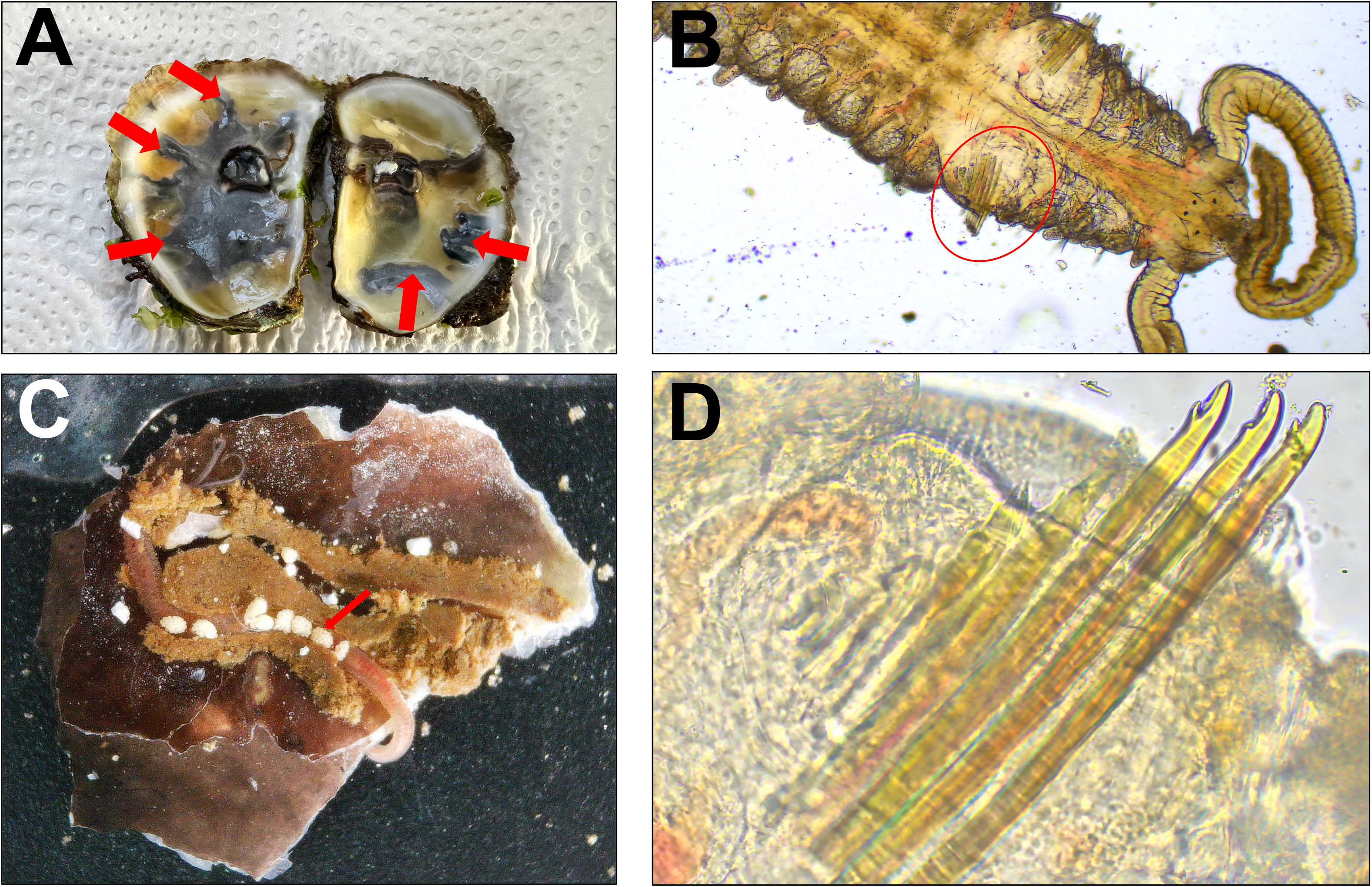
*Polydora websteri* infesting oysters from Cape Cod. A. Extensive mud blisters on the inner layer of the shell (arrows), B. Partial dorsal view of *P. websteri* showing its characteristic modified fifth chaetiger spines (circle), C. Female worm brooding her egg string in an exposed burrow, D. Close-up view of *P. websteri*’s modified fifth chaetiger spines (companion chaetae not shown).

**Fig. 6.**
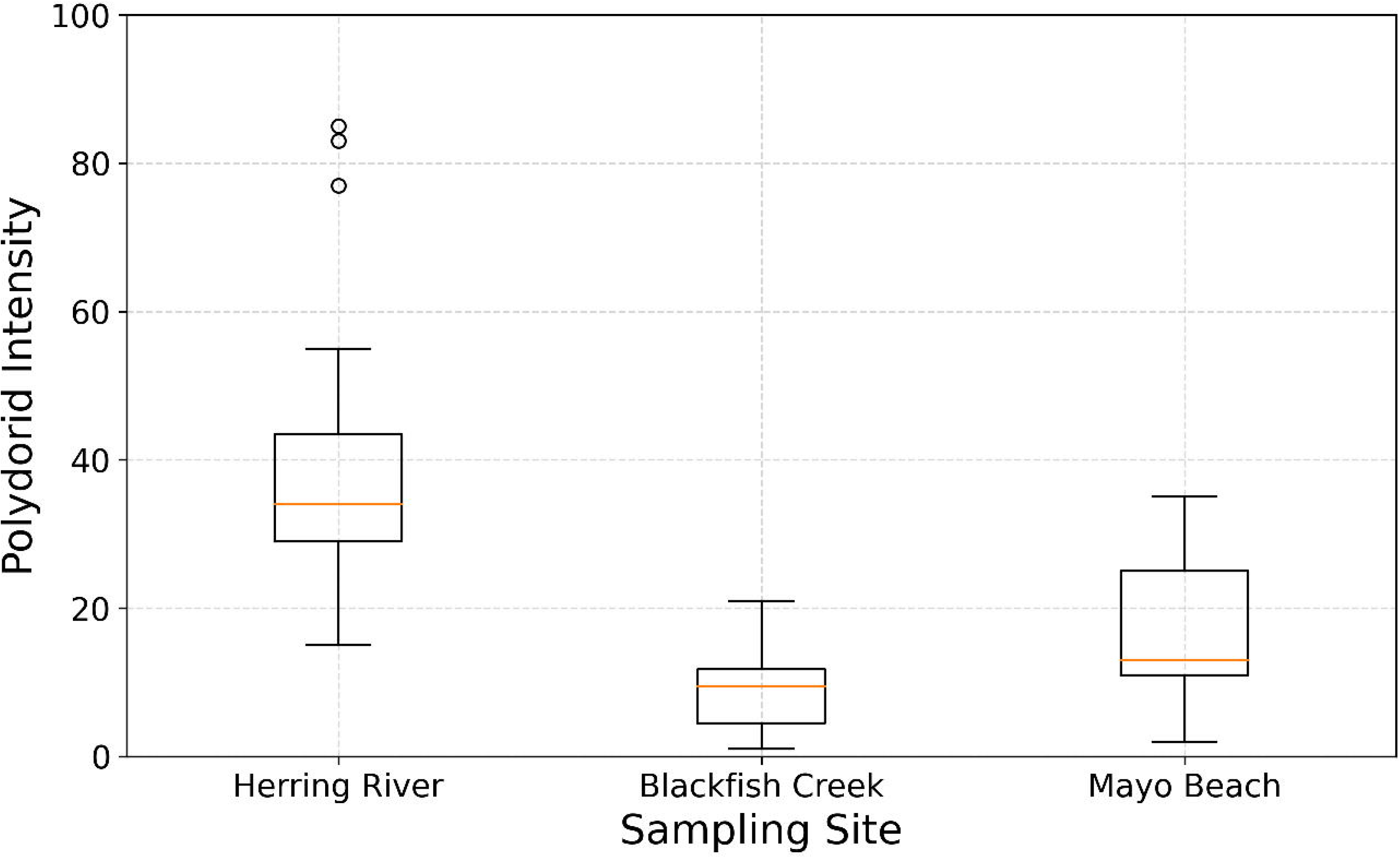
Boxplot showing mean intensity of *Polydora websteri* in oysters sampled from three sites in Wellfleet Harbor, Massachusetts.

**Fig. 7.**
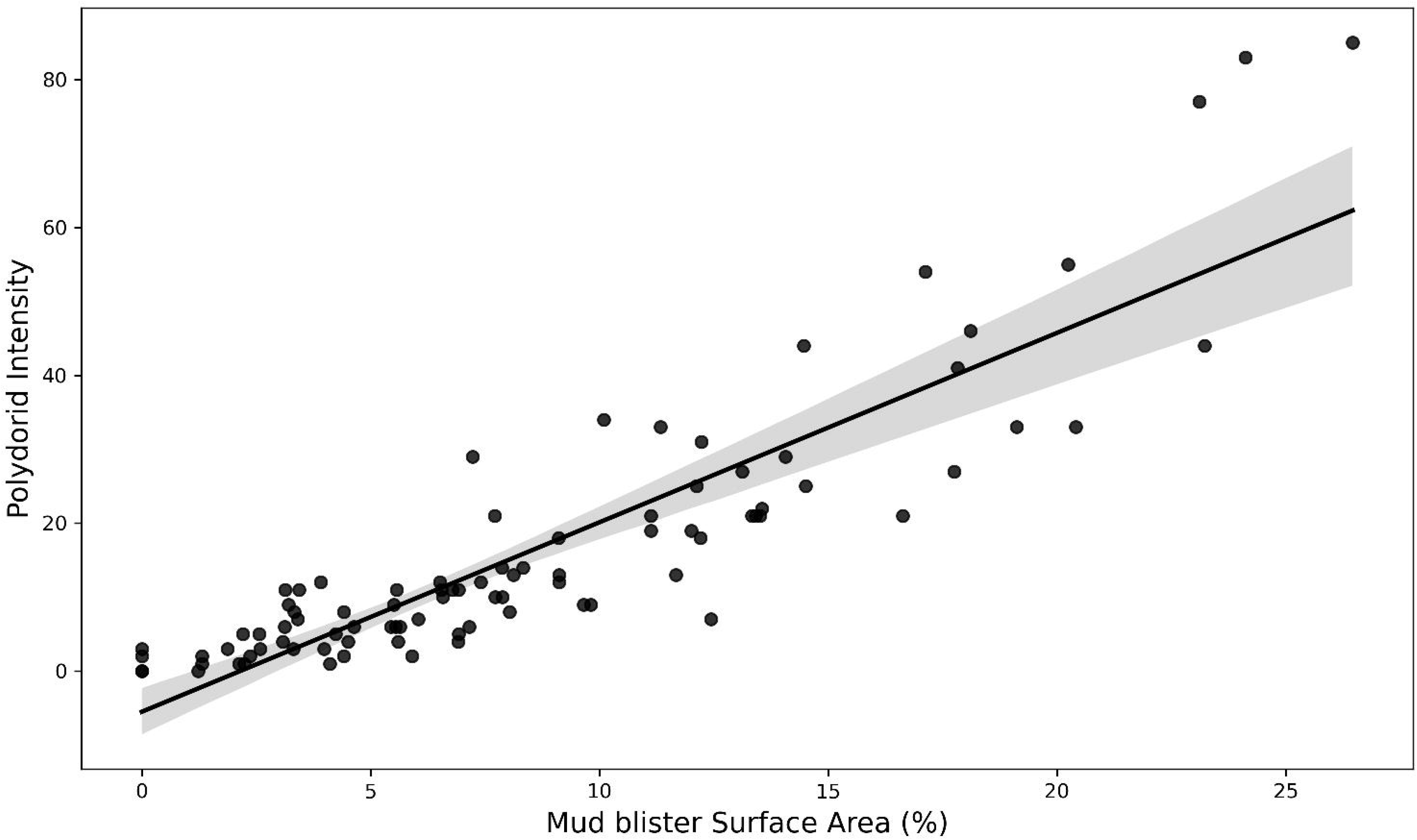
Linear regression showing relationship between mud-blister surface area and polydorid intensity. Shaded region represents 95% confidence bands.

## 4. Discussion

The purpose of this study was to evaluate parasite load, identity and distribution of shell-boring polychaetes associated with shellfish collected from wild and farmed populations in an important oyster culturing region of Cape Cod, Massachusetts. In summary, the major findings showed that oyster populations from all three sites in Wellfleet Harbor were infested by *Polydora websteri* although infestation rates in the Herring River oysters were far higher than those from Black Fish Creek and Mayo Beach. In addition, *P. websteri* from these sites showed high levels of genetic admixture with not only cohorts from other New England regions but also with other globally distant populations.

Considering that all three sampling sites are located within the same harbor in close proximity to each other (< 10 km apart) and the fact that all sampled oysters were within the same size range, it was surprising to observe significant discrepancies in infestation levels. One explanation lies in the nature of the Herring River site, which for more than a century has been cut off from normal tidal flow due to the presence of a dike constructed in 1909 for mosquito eradication and development of arable land (Portnoy and Allen, 2006). Such a fragmented system eliminates tidal flushing, increases residence time and thus reduces water flow and subsequent exchange with Cape Cod Bay. Though they did not explicitly test the effects of tidal flow, Cole et al. (2020) suggested that tidal excursions may transport *P. websteri* larvae away from a farmed site. This means that for a site like the Herring River, larvae can recruit easier to their natal sites without being swept away by tidal currents, resulting in a massive aggregation of worms within a relatively small area. Handley and Bergquist (1997) also found that intertidal exposure or tidal currents were important in mediating infestation levels of polydorid worms. Aside from the lack of tidal flow, high fecal coliform bacteria levels have been consistently reported from the Herring River, which consequentially has resulted in the closure of shellfish beds for both commercial and recreational harvesting (Portnoy and Allen, 2006). Whether high bacterial levels are related to the increased proliferation of shell-boring *Polydora* worms is currently not known but is an area ripe for future investigation.

While previous studies have quantified the relationship between *Polydora* infestation and oyster size and condition, this is the first study to directly assess the relationship between mud-blister area and polydorid infestation level. A linear regression analysis of the pooled data showed that the size of mud-blisters, specifically their surface area, is a reliable predictor of *Polydora* intensity, at least with respects to *P. websteri*. Working with restaurant stakeholders, this can be useful for shellfish growers to determine the state of their crop after a harvest. For example, restaurants who purchase oysters from a particular grower could take pictures of the recycled shells which growers can then use to estimate infestation levels on their oyster farm.

DNA barcoding analysis unequivocally confirmed that *Polydora websteri* is the sole culprit responsible for infestation of Wellfleet oysters. The genetic analyses also found low levels of genetic structuring between worms from Wellfeet and other local, regional and global populations. Even with the addition of *P. websteri* sequences from two new geographic regions in Eastern Europe (Black Sea and Sea of Azov) along with additional sites sampled in this study from New England, our results support recent findings by Rice et al. (2018) and Rodewald et al. (2021) who showed that *P. websteri* is a genetically homogenous species. These authors attributed the high levels of admixture to the aquaculture trade where historical and current movement of infested oysters both regionally and internationally has resulted in high levels of gene-flow among *P. websteri* populations, some of which would otherwise have been isolated. Indeed, the haplotype network generated in this study which shows a highly mixed common haplotype combined with a very small number of mutational steps, is indicative of panmixia (David, 2018). In such a scenario, multiple and repeated introductions have diluted the effects of phylogeographic barriers, rendering them virtually non-existent (David, 2018; David and Loveday, 2018). The Wellfleet worms showed the lowest levels of genetic structuring (highest levels of connectivity) with worms sampled from *C. virginica* in Maine (Φ_ST_ = 0.013) which is not surprising considering the long legacy of oyster transplantation along the east coast of the United States (Brennessel, 2008). However, the low level of genetic structure observed between Wellfleet specimens and those from eastern Europe (Φ_ST_ = 0.058) was surprising. The eastern European *P. websteri* were sampled from two different hosts (*C. gigas* and the ark clam *Anadara kagoshimensis*) in the Black Sea and the Sea of Azov respectively – both of which are isolated from the western Atlantic. *Crassostrea gigas* was first intentionally introduced to the Black Sea in the 1980s from Japan (Zolotarev, 1996) for aquaculture purposes and has since extended its range to form ‘naturalized’ populations in this region (Aydin and Mustafa, 2021). *Anadara kagoshimensis* is regarded as a ‘self-introduced’ species to the Sea of Azov where it has naturally expanded its range over the past few decades from the Black Sea basin (Zhivoglyadova et al. 2021). However, it was only recently that *P. websteri* was found burrowing into *A. kagoshimensis* in this region (Syomin et al. 2021). Interestingly, *P. websteri* and *A. kagoshimensis* have overlapping ranges in the nearshore waters of Japan indicating that *P. websteri*’s introduction to both the Black and Azov Seas may have been due to the anthropogenic movement of bivalves from one or multiple waterbodies in Asia. In a similar manner, Rice et al. (2018) also hypothesized that *P. websteri* was likely introduced to the United States from Asia with the latter being the species’ native range based on genetic diversity estimates. However, we should note that the true native range of *P. websteri* cannot be definitively resolved without a more comprehensive sequence dataset. Regardless of where *P. websteri*’s native range lies, it is clear that a complex history of bivalve introductions – both natural and anthropogenic, has resulted in unusually high levels of genetic connectivity among the worm’s populations around the globe. Today, New England and much of the United States have stricter biosecurity controls than in the past and so it is very unlikely that the low levels of genetic differentiation observed between U.S. specimens and other distant populations like the Black Sea worms are due to recent introductions. Instead, it is more likely that the current mtDNA genetic patterns is reflective of deep historical dispersal. We suggest that future studies on *P. websteri* deploy more sensitive and high-resolution markers such as RAD-Seq which may uncover finer scale connectivity patterns for this species.

### 4.3. Conclusions and implications for oyster farming in Cape Cod

In 2022, the National Oceanic and Atmospheric Administration (NOAA) announced a 5-year strategic plan to enhance the growth of sustainable aquaculture in the United States (NOAA Aquaculture Strategic Plan). Goal two of this plan emphasizes sustainability which involves “advancing aquatic health management practices” along with understanding “climate change effects”. Shell-boring polydorids pose a direct threat to the former while the latter may potentially interact with polydorid life history, resulting in a negative synergistic effect that greatly increases the risk of shellfish extirpation in the US (David, 2021). Unfortunately, attempts to completely eradicate polydorids from shellfish farms once they have been infested have not yet been demonstrated in the literature. Currently, it appears that reducing prevalence and intensity levels may be the only feasible option. Two recent literature reviews by Spencer et al. (2021) and Tan et al. (2023) summarized treatments that have shown to be somewhat successful at accomplishing this. These include hyper- and hypo saline (freshwater shock) treatments, air drying at low temperatures, and shell coating. For shellfish farms in the Cape such as the ones sampled in this study, oysters are sometimes stored in dry cellars during periods of extreme cold – a practice known as ‘pitting’ (Brennessel 2008). This may explain why prevalence and intensities were significantly lower in the MB and BFC sites compared to the Herring River where the oysters are almost never exposed to air. In addition to promoting this air-drying procedure (which is incidental due to the climate that these oysters are cultured in), hyposaline treatments may also be a potential option for farmers, especially during the critical months before harvesting when water temperature increases – a condition that favors polydorid reproduction and growth (David and Simon, 2014).

As of 2023, the town of Wellfleet is engaged in a restoration project which will involve removal of the dike to restore tidal flow with the hopes of reopening shellfish beds for harvesting. While polydorids are not a health hazard for humans, heavy infestation in these oysters renders them commercially inviable. While recreational and subsistence shellfish harvesters might tolerate *Polydora-*riddled oysters, those who are involved in the commercial aquaculture trade may not find profitability in these oysters since their marketability has been compromised due to extensive mud-blisters. It is therefore recommended that additional surveys to this one be implemented in the Herring River sometime after the removal of the dike to assess any temporal changes in polydorid composition and infestation levels.

## Acknowledgements

We would like to thank the oyster farmers at Black Fish Creek and Mayo Beach in the Town of Wellfleet for providing access to their oyster farms.

